# *N*-hydroxy pipecolic acid methyl ester is involved in Arabidopsis immunity

**DOI:** 10.1101/2022.06.03.494702

**Authors:** Lennart Mohnike, Weijie Huang, Brigitte Worbs, Kirstin Feussner, Yuelin Zhang, Ivo Feussner

## Abstract

The biosynthesis of *N*-hydroxy pipecolic acid (NHP) has been intensively studied, though knowledge on its metabolic turnover is still scarce. To close this gap, we discovered three novel metabolites via metabolite fingerprinting in *Arabidopsis thaliana* leaves. Exact mass information and fragmentation by mass spectrometry (MSMS) suggest a methylated derivative of NHP (MeNHP), a NHP-*O*Glc-hexosyl conjugate (NHP-*O*Glc-Hex) and an additional NHP-*O*Glc-derivative. All three compounds were formed in wildtype leaves but not present in the NHP deficient mutant *fmo1-1*. The identification of these novel NHP-based molecules was possible by a dual-infiltration experiment using a mixture of authentic NHP- and D_9_-NHP-standards for leaf infiltration followed by an UV-C treatment. Interestingly, the signal intensity of MeNHP and other NHP-derived metabolites increased in *ugt76b1-1* mutant plants. This suggests a detour, for the inability to synthesize NHP-*O*-glucoside. For MeNHP, we unequivocally determined the site of methylation at the carboxylic acid function. MeNHP application by leaf infiltration leads to the detection of a MeNHP-*O*Glc as well as NHP, suggesting MeNHP-hydrolysis to NHP. This is in line with the observation that MeNHP-infiltration is able to rescue the *fmo1-1* susceptible phenotype against *Hyaloperonospora arabidopsidis* Noco 2. Together these data suggest MeNHP as additional storage or transport form of NHP.

**Highlight:** In this work, we identify *N*-hydroxy pipecolic acid (NHP) metabolites including methyl ester and complex glycosides. The application of methyl ester is able to rescue the disease phenotype of the biosynthesis deficient mutant of NHP.

## Introduction

Plants encounter reduced growth or induce early senescence, if they are unable to maintain a balance between growth and defense (von Saint Paul *et al*., 2011; Zhang *et al*., 2017). Their immune system depends on a tightly regulated and highly dynamic balance of activation and inactivation (Karasov *et al*., 2017; Zeier, 2021). Salicylic acid (SA) and *N*-hydroxy pipecolic acid (NHP) are two key molecules to concert the defense response against (hemi-)biotrophic pathogens (Fu and Dong, 2013; Hartmann and Zeier, 2019).

In the Brassicaceae model organism *Arabidopsis thaliana*, SA and NHP are synthesized upon pathogen infection. Roughly, 90 percent of SA derive from chorismic acid, which is converted via the ISOCHORISMIC ACID SYNTHASE 1 (ICS1) pathway. This pathway features the enzymes AvrPphB SUSCEPTIBLE 3 (PBS3) and ENHANCED PSEUDOMONAS SUSCEPTIBILITY 1 (EPS1) to synthesize SA (Rekhter *et al*., 2019; Torrens-Spence *et al*., 2019; Wildermuth *et al*., 2001). NHP derives from L-lysine via the enzymatic route of AGD2-LIKE DEFENSE RESPONSE PROTEIN 1 (ALD1), SYSTEMIC ACQUIRED RESISTANCE DEFICIENT 4 (SARD4) and FLAVINE-DEPENDENT MONOOXYGENASE 1 (FMO1) (Fig. 1). Both molecules orchestrate defense signaling including the activation of protective measures, such as defense gene expression, and danger signal amplification (Chen *et al*., 2018; Ding *et al*., 2016; Hartmann *et al*., 2018; Navarova *et al*., 2012). In consequence, to prime distant leaf tissue for robust defense against secondary stressors, termed systemic acquired resistance (SAR) (Chen *et al*., 2018; Hartmann *et al*., 2018).

**Fig. 1.**
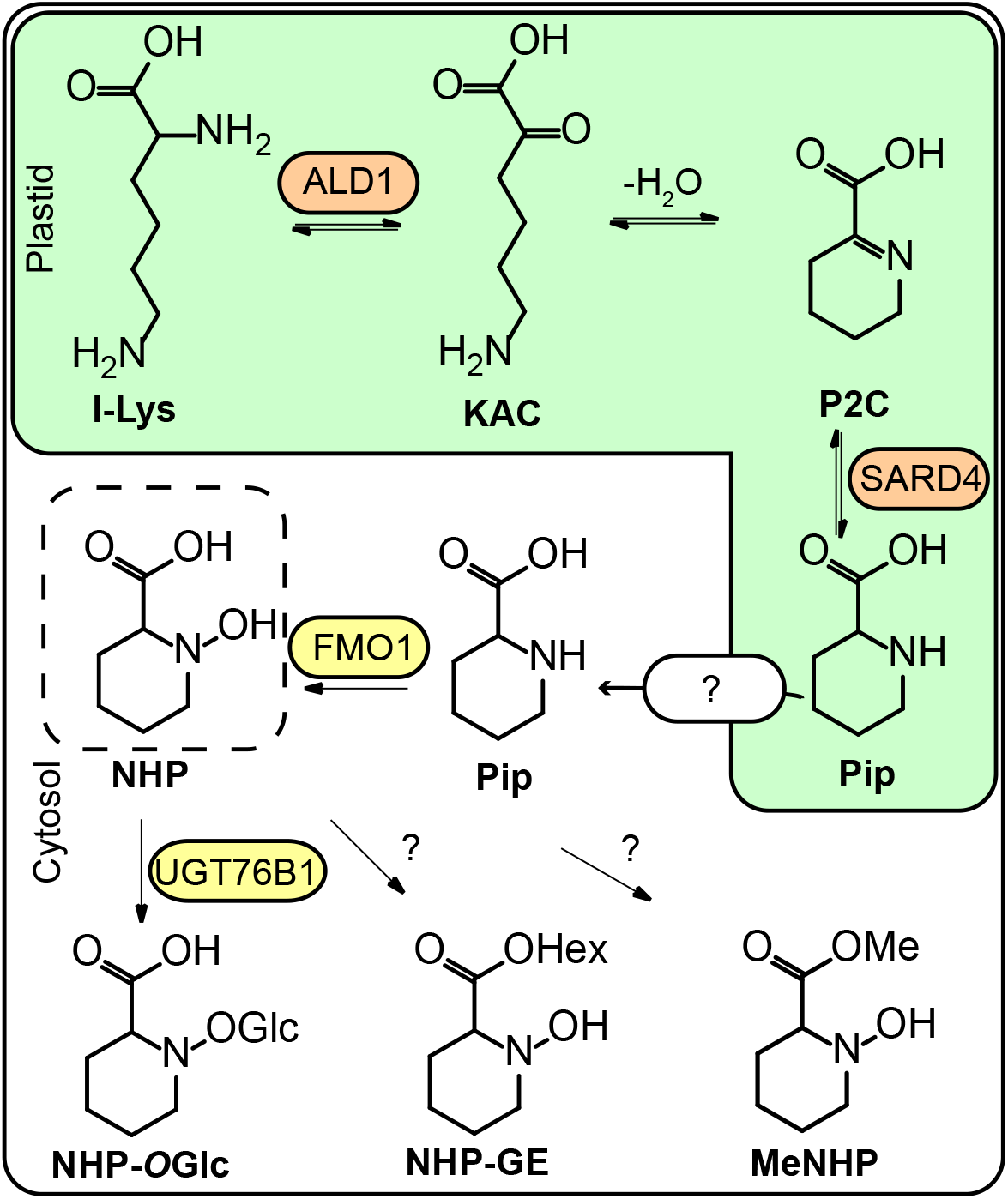
Biosynthesis route of NHP-metabolites in *Arabidopsis*. In the plastid, l-lysine (l-Lys) is converted by AGD2-LIKE DEFENSE RESPONSE PROTEIN 1 (ALD1) to epsilon-amino-alpha-keto caproic acid (KAC). Via spontaneous cyclization under water loss of KAC, piperidein-2-carboxylic acid (P2C) is formed. A reductase capable to reduce P2C to pipecolic acid (Pip) is SYSTEMIC ACQUIRED RESISTANCE DEFICIENT 4 (SARD4). How pipecolic acid is exported from the chloroplast is still elusive. The FLAVINE-DEPENDENT MONOOXYGENASE 1 (FMO1) catalyzes *N*-hydroxylation of Pip, resulting in *N*-hydroxy pipecolic acid (NHP) (Chen *et al*., 2018; Ding *et al*., 2016; Hartmann *et al*., 2018; Navarova *et al*., 2012). NHP was shown to be glycosylated by UGT76B1 to NHP-*O*-glycoside (NHP-*O*-Glc). Furthermore, NHP-glycoside-ester (NHP-GE) was described, but respective enzyme is not known (Bauer *et al*., 2021; Hartmann and Zeier, 2018). Methylation of NHP to NHP-methyl-ester (MeNHP) was shown as an additional mechanism of NHP-turnover *in planta* in this study.

Abnormal accumulation of plant defense hormones can lead to phenotypes such as “dwarfism” or early senescence (Cai *et al*., 2021; von Saint Paul *et al*., 2011; Zhang *et al*., 2017). One way to regulate the cellular concentrations of SA and NHP is by metabolic turnover. SA is glycosylated by a minimum of three UDP-dependent glycosyltransferases (UGTs): UGT74F1, UGT74F2 and UGT76B1, forming the SA-glycoside (SAG) (Dean and Delaney, 2008; George Thompson *et al*., 2017; Maksym *et al*., 2018; Noutoshi *et al*., 2012). In addition, UGT74F2 was shown to produce SA-glycoside-ester (SGE) (Dean and Delaney, 2008; George Thompson *et al*., 2017). Another mechanism of SA turnover is via the 3- and 5-hydroxylation via SA-3-hydroxylase (S3H) and SA-5-hydroxylase (S5H) (Zhang *et al*., 2013; Zhang *et al*., 2017). The metabolic products of the reactions are 2,3-di-hydroxy benzoic acid (2,3-DHBA) and 2,5-di-hydroxy benzoic acid (2,5-DHBA). These molecules themselves can be turned-over by UGT76D1 to 2,3- and 2,5-DHBA-glycosides (2,3- and 2,5-DHBA), respectively (Huang *et al*., 2018). Arabidopsis plants harboring a mutation in *S5H* exhibit reduced growth and increased defense responses. The *s3h s5h* double mutant shows further increase in SA levels, reduced growth and enhanced resistance compared to wild type plants (Zhang *et al*., 2017).

The identification of the *S*-adenosyl-dependent methyl transferase BENZOIC ACID/SA METHYL-TRANSFERASE 1 (BSMT1) and its volatile aromatic-ester product SA methyl ester (MeSA) opened novel aspects of defense priming and SAR (Chen *et al*., 2003; Park *et al*., 2007). MeSA has been shown to allow defense priming in systemic leaves and airborne plant-to-plant communication resulting in acquired immunity in receiver plants (Park *et al*., 2007; Shulaev *et al*., 1997). Nevertheless, the function of MeSA in Arabidopsis SAR is still under discussion. The Arabidopsis *bsmt1* mutant plants exhibit a wild type-like SAR response, without significant accumulation of MeSA in response to pathogen infection (Attaran *et al*., 2009). Additionally, *bsmt1* mutants are not compromised in communicating airborne SAR induction (Wenig *et al*., 2019). MeSA can be metabolized to MeSAGlc by UGT73C1 in Arabidopsis (Chen *et al*., 2019).

In terms of the SAR mediator NHP, only two products of turnover were described. NHP glucosylation was identified in several independent studies resulting in the formation of NHP-*O*Glc (Bauer *et al*., 2021; Cai *et al*., 2021; Holmes *et al*., 2021; Mohnike *et al*., 2021). Bauer and colleagues have proposed a second glycoside form, NHP-Glc-ester (NHP-GE) (Bauer *et al*., 2021). Nevertheless, the identification of NHP metabolites may so far be incomplete. Other modifications, such as, methylation or amino acid conjugation were not described for NHP yet.

Here we report infection and UV-C-dependent formation of methylated NHP (MeNHP) identified via ultra-high-performance-liquid-chromatography high-resolution mass spectrometry (UHPLC-HRMS) metabolome analysis. We confirmed NHP-methylation via D_9_-labeled NHP and determined carboxylic acid methylation via a comparison to a synthesized authentic MeNHP-standard. Furthermore, we showed that MeNHP is able to rescue the NHP-deficient phenotype of *fmo1-1* mutant plants and reduce oomycete spore growth in *Arabidopsis thaliana* interaction. In addition, we present a dual-infiltration experiment of a mixture of NHP and D_9_-NHP, to identify and to investigate novel metabolites of NHP in a non-targeted manner.

## Material and methods

### Plant material and growth conditions

*Arabidopsis thaliana* ecotype Col-0, *fmo1-1, ugt76b1-1, fmo1 ugt76b1* (CRISPR *ugt76b1-5* in *fmo1-1*) and *FMO1-3D* were used in this study accordingly (Mohnike *et al*., 2021). Additionally, *FMO1-3D ald1* was used in this study. We obtained *nhpmt1-1* (SALK_053006) and *nhpmt1-2* (SALKseq_135601) mutant plants from SALK Nottingham. Plants were grown on steam sterilized soil under short day (8h light/16h dark) or long day (16h light/8h dark) for 4 and 6 weeks. The light intensity was 100-120 µmol/m^2^/s and the humidity was 80 % relative unless specified. The light source were MASTER LED tubes HF 600mm HO 8W840 T8 (PHILIPS AG, Amsterdam, Netherlands).

### Pseudomonas infection and UV-treatment

To induce defense metabolism, plants were treated with *Pseudomonas* bacteria or UV-C light. *Pseudomonas syringae* strain ES4326 (*P*.*s*.*m*.) were grown in LB-medium with 25µg Rifampecin overnight at 28 °C. The culture was pelleted, medium was decanted, and the bacteria were suspended in 10 mM MgCl_2_. Bacteria were diluted to OD_600_=0.05 and infiltrated to the abaxial side of the leaf. As mock treatment control 10 mM MgCl_2_ were infiltrated to the leaf. Plants were incubated for 8, 24, or 48 hours, as stated accordingly in the results section and figure legends. UV-C treatment was performed for 20 min in a PrettleTelstar sterile bench as described (Mohnike *et al*., 2021). Plants were incubated for 24 hours post treatment if not stated otherwise.

### Chemical synthesis of MeNHP

MeNHP was synthesized from methylpipecolinate hydrochloride after a modified procedure (de Sousa 2016, Frontiers). See supplemental material for a detailed description of the synthesis route, materials and techniques used to gain MeNHP. Additionally we deposited NMR-spectra and MS-spectra of the quality control measures.

### Dual infiltration of authentic NHP and D_9_-NHP standard

Both 1 mM NHP and D_9_-NHP, respectively, in 10 mM MgCl_2_ were co-infiltrated to 3 leaves of each individual plant of Col-0, *fmo1-1* and *ugt76b1-1*. As mock treatment 10 mM MgCl_2_ was infiltrated accordingly. Both mock and NHPs infiltrated plants were either kept further untreated or were exposed to UV-C radiation for 20 minutes, as described above. Plants were incubated for 24 hours post treatment. Samples were harvested and stored in -80 °C until extraction with 80 %MeOH.

### MeNHP infiltration for metabolite tracking

1 mM MeNHP were directly infiltrated to the apical side of the leaf. MeNHP was solved in 10 mM MgCl_2_. The infiltrated plants were incubated for 24 hours. Leaves were harvested and frozen in liquid nitrogen. The samples were stored at -80 °C.

### MeNHP induced resistance assay

To investigate the MeNHP induced resistance, ddH_2_O (mock) or MeNHP at the indicated concentrations diluted in ddH_2_O were infiltrated with a needleless syringe on two full-grown leaves of 3-week-old *fmo1-1* and Col-0 plants. 24 hours post infiltration, plants were challenged with *Hyaloperonospora arabidopsidis* Noco 2 by spraying a conidiaspore solution at a concentration of 50,000 spores/mL. The challenged plants were then grown in a plant chamber at 18°C with a relative humidity of 80% under short day cycle (8-h light/16-h dark). Infection was scored at 7 days post inoculation as described previously (Ding *et al*. 2016). In brief, infection was scored by the conidiaspore growth on distal leaves with the following rating category: Category 5 = more than 5 conidiaspores observed on more than 2 distal leaves, Category 4 = more than 5 conidiaspores observed on 2 distal leaves, Category 3 = less than 5 conidiaspores observed on 2 distal leaves, Category 2 = more than 5 conidiaspores observed on 1 distal leaf, Category 1 = less than 5 conidiaspores observed on 1 distal leaf, Category 0 = none conidiaspore observed on all distal leaves.

### Extraction of plant metabolites

Metabolite extracts were generated from frozen leaf material. Leaves were ground under liquid nitrogen and weight to 100 mg fresh weight (MTBE, only Fig. 2) or 50 mg (80 % MeOH). The MTBE extraction was performed as described earlier (Mohnike *et al*., 2021). The 80 % MeOH extraction was slightly modified of an extraction protocol (kindly communicated by Prof. Dr. Armin Djamei). 50 mg ground leaf material were given into 2 mL Eppendorf cups and 800 µL of 80% MeOH were added. The samples were vortexed to secure homogenization. Afterwards, ultrasonication was applied to the samples two times for each 15 min. The samples were centrifuged at 18.000 ×g for 15 min. 700 µL of debris free supernatant were transferred into new tubes and evaporated under streaming nitrogen. Metabolites were resolved in 20 % MeOH by vortex. The solutions were centrifuged at 18.000 ×g for 10 min prior to LC-analysis to remove remaining debris. 80 µL were transferred into the LC-MS vials.

**Fig. 2.**
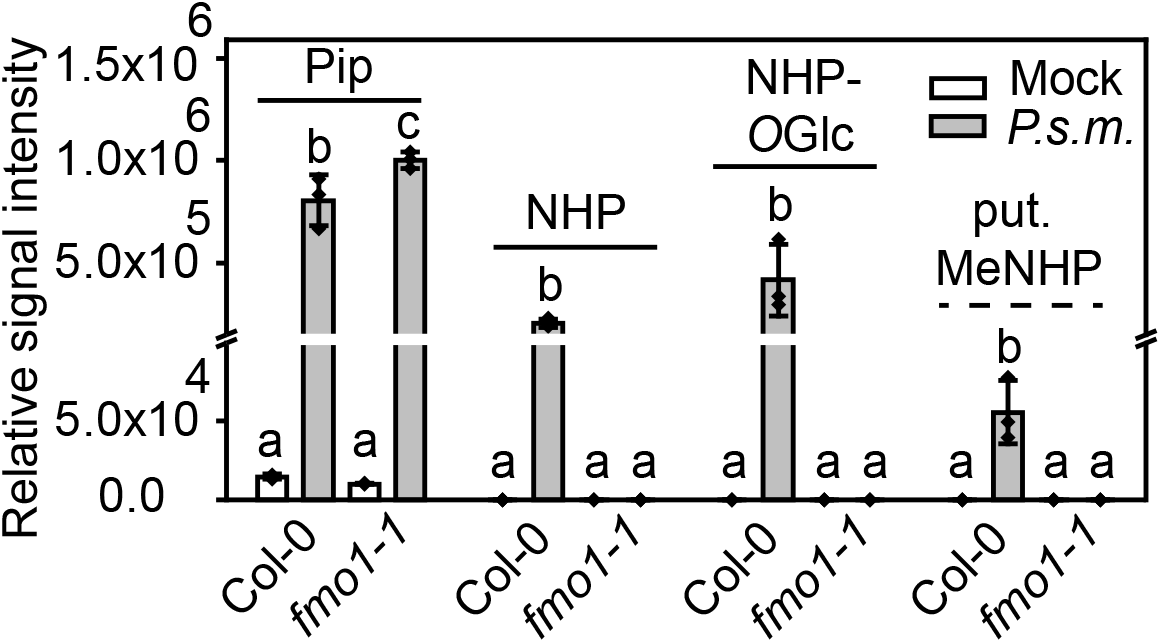
Accumulation of selected *FMO1*-dependent and independent metabolites in Col-0 and *fmo1-1* leaves in response to *P*.*s*.*m*. infiltration. *A. thaliana* plants grown for 6 weeks under short-day conditions (8h light/16h dark) were infiltrated with 10 mM MgCl_2_ (Mock) or virulent *P. syringae* ES4326 strain at OD_600_= 0.05 in 10 mM MgCl_2_ (*P*.*s*.*m*.). Infiltrated leaves were harvested 24 hours post infiltration, extracted by MTBE-procedure and analyzed via UHPLC-HRMS. Data were processed using Profinder 8.0 (Agilent Technologies, Santa Clara, CA, USA) and OriginPro2020 (Origin, San Francisco, CA, USA). Relative signal intensities of Pip, NHP, NHP-*O*Glc and putative methyl-NHP (put. MeNHP) are shown. Y-axis has a brake between 1×10^5^ and 1.5×10^5^. Bars represent mean values with standard deviation of n=3 replicates. Letters indicate statistical differences, individually calculated for each metabolite (p < 0.05, one-way ANOVA post-hoc tukey-test). Each replicate represents an independent pool of nine infiltrated leaves from three plants.

### UHPLC-HRMS-based metabolite fingerprinting

Metabolite fingerprinting was conducted according to Feussner and Feussner 2019, as described in Mohnike et al. 2021 (Feussner and Feussner, 2019; Mohnike *et al*., 2021). In brief, extracted samples were analyzed with the UHPLC1290 (Agilent Technologies, Santa Clara, CA, USA) coupled to a HRMS instrument 6540 UHD Accurate Mass Q-TOF (Agilent Technologies) with Agilent Dual Jet Stream Technology as electrospray ionization (ESI) source (Agilent Technologies). The ACQUITY HSS T3 column (2.1 × 100 mm, 1.8 µm particle size, Waters Corporation) was used for chromatographic separation at a flow rate of 500 µL/min at 40 °C. The solvent system applied A (water, 0.1 % (v/v) formic acid) and B (acetronitrile, 0.1 % (v/v) formic acid) were used. The gradient applied was: 0 to 3 min 1 % to 20 % B; 3 to 8 min 20 %to 97 % B; 8 to 12 min: 100 % B; 12 to 15 min: 1 % B. For technical details were described recently (Mohnike et al. 2021). Data were acquired using Mass Hunter Acquisition B.03.01. Data deconvolution was performed using Profinder 10.0 (Agilent Technologies). Data were processed using MarVis-Suite (Kaever *et al*., 2015; Kaever *et al*., 2012; Kaever *et al*., 2009) (http://marvis.gobics.de) or OriginPro2020 (OriginLab Corporation, Northampton, MA, USA).

## Results

### Identification of MeNHP via metabolite fingerprinting

Following the hypothesis that molecules of NHP-turnover are missing in *fmo1-1* plants, we compared Col-0 wild type against *fmo1-1* leaves that were infected with *P*.*s*.*m*. The leaf extracts were analyzed via UHPLC-HRMS and the obtained dataset was searched for hypothetical NHP-metabolites based on previously described modifications of SA and JA, for instance, hydroxylation, dehydrogenation, decarboxylation and methylation. As proof of concept, the dataset was analyzed for Pip, NHP and NHP-*O*Glc accumulation after *P*.*s*.*m*. treatment. As expected, NHP and NHP-*O*Glc were not detected in *fmo1-1* plants (Fig. 2). However, we detected exclusively in Col-0 a relative signal intensity with a mass-to-charge ratio (*m*/*z*) of 160.097 in the positive ionization mode and a retention time of 2.63 minutes, which corresponds to the mass of methylated-NHP (put. MeNHP). The exact mass of this molecule has been calculated with 159.090 Da (C_7_H_13_NO_3_), showing a mass shift of 14.015 Da to NHP. This shift is equivalent to a methyl group deriving possibly from methylation of NHP.

### MeNHP is a metabolite of NHP and its identity was unequivocally confirmed by authentic NHP-methyl-ester-standard

To confirm MeNHP as a metabolite of NHP, therefore, being dependent on functional *FMO1*, we infiltrated labeled D_9_-NHP into Col-0 and *fmo1-1* leaves and measured formation of D_9_-MeNHP. Indeed, we detected D_9_-MeNHP in both Col-0 and *fmo1-1* leaves. The labeled compound had a retention time shift towards a polar elution compared to MeNHP. Non-labeled native MeNHP was again only found in Col-0 leaves (Fig. 3A).

**Fig. 3.**
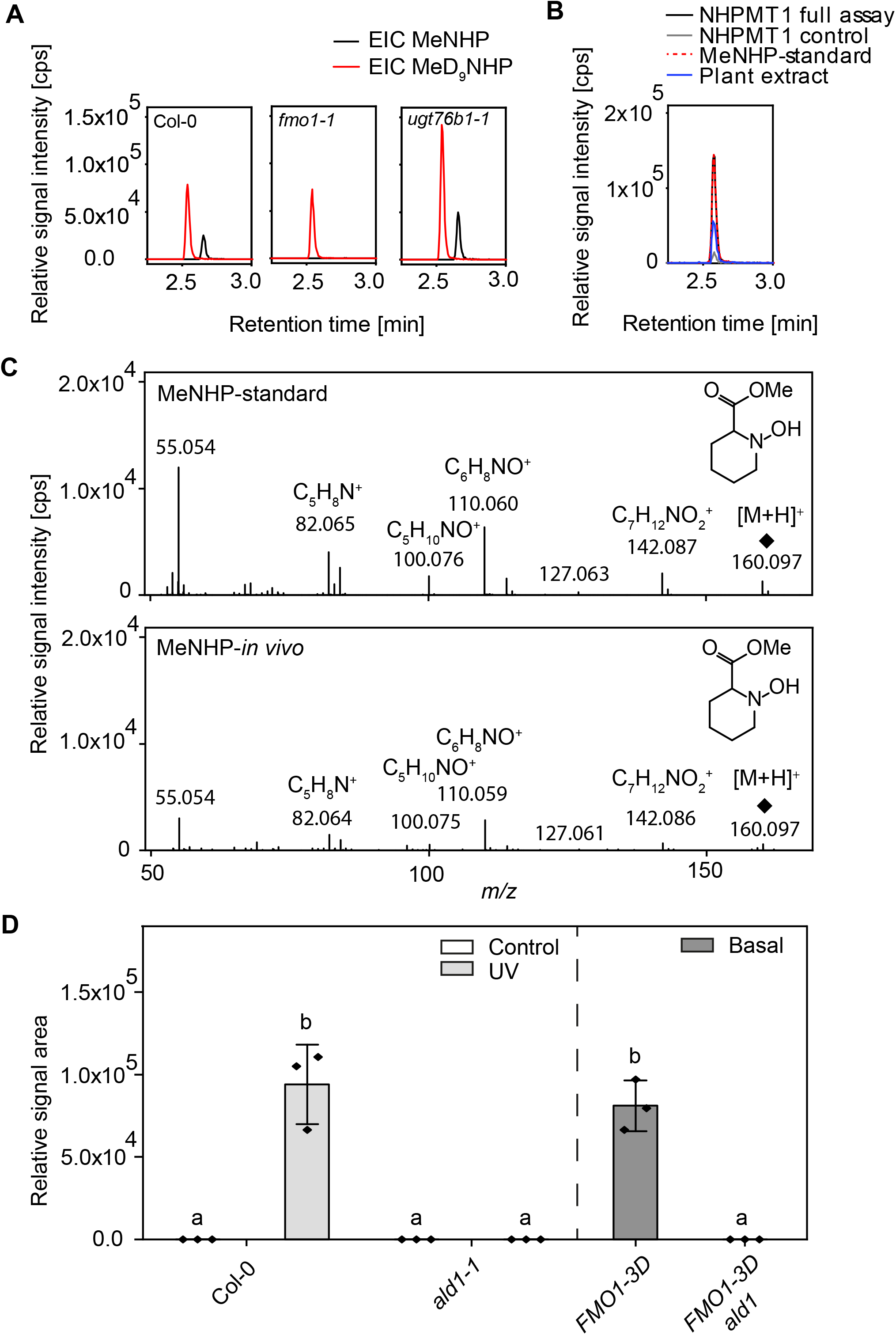
Characterization of *N*-hydroxy pipecolic acid methyl ester (MeNHP). **A** Extracted ion chromatograms of MeNHP [M+H]^+^ 160.097 and D_9_-MeNHP [M+H]^+^ 169.153 in Col-0, *fmo1-1* and *ugt76b1-1* leaves. Col-0, *fmo1-1* and *ugt76b1-1* leaves were infiltrated with D_9_-NHP. After 24 hours, leaves were harvested, extracted and analyzed by UHPLC-HRMS. **B** Extracted ion chromatograms at [M+H]^+^ 160.097 of *in vivo* metabolite extract, synthesized authentic NHP-methyl-ester standard, *in vitro* assay with NHPMT1 and *in vitro* assay with boiled NHPMT1 as control. **C** Comparison of MSMS-fragments of authentic MeNHP-standard and *in planta* MeNHP. **D** Relative amount of MeNHP in Col-0, *ald1, FMO1-3D* and *FMO1-3D ald1* plants. Col-0 and *ald1* plants were kept untreated or treated with UV-C for 20 min. Plants were incubated for 24 hours before harvesting. *FMO1-3D* and *FMO1-3D ald1* plants were left untreated before harvesting. Data represents mean of three biological replicates for the MeNHP signal (relative signal area) via UHPLC-HRMS analysis. Error bars represent standard deviation. Letters indicate statistical differences, individually calculated for each experiment (p < 0.05, one-way ANOVA post-hoc tukey-test).

Next, we developed a strategy for the chemical synthesis of a NHP-methyl-ester standard to confirm the identity of MeNHP unequivocally. NHP-methyl-ester was synthesized from methyl piperidin (Supplementary File). Additionally, we tried to identify potential methyl transferase candidates with publicly available co-expression data files of the NHP-metabolizing enzyme UGT76B1 (ATTED II, version 11.0). One gene of interest was AT4G22530 (put. NHP-methyl transferase 1 (NHPMT1) which was annotated as *S*-adenosyl methionine-dependent methyl transferase and its expression is NHP-responsive (Yildiz *et al*., 2021). The cDNA of the gene was cloned into a pET28a-expression vector, and the encoded protein was heterologously expressed in *E. coli*, purified to homogeneity, used for an *in vitro* activity assay with NHP as substrate. The reaction was followed by UHPLC-HRMS (Fig. S1). Indeed, the authentic MeNHP-standard coeluted with *in planta* and enzymatically generated MeNHP at a retention time of 2.57 min and is presented in the extracted ion chromatogram of *m/z* 160.097 (Fig. 3B).

In addition, the fragmentation pattern of the MS/MS-spectra of *m/z* 160.097 (MeNHP) exhibit identical fragments as the *in vivo* derived MeNHP and the authentic standard. The main fragments are *m/z* 142.08, *m/z* 127.063, *m/z* 110.060, *m/z* 100.076 and *m/z* 82.065 (mass accuracy of ± 2 mDa) (Fig. 3C). The fragment *m/z* 142.08 represents C_7_H_12_NO_2_^+^ after a loss of the *N*-hydroxy function, comparable to NHPs-fragment *m/z* 127.063 (Fig. S2). Moreover, identical fragmentation behavior of MeNHP (C_7_H_13_NO_3_) and NHP (C_6_H_11_NO_3_) is observed by fragment ions *m/z* 110.06, *m/z* 100.07 and *m/z* 82.06. First, *m/z* 110.06 represents C_6_H_8_NO^+^ after loss of two hydroxy functions. Second, *m/z* 100.07 represents the fragment C_5_H_10_NO^+^ obtained by the loss of the carboxylic acid function with NHP and methyl carboxylic acid function with MeNHP. Last, *m/z* 82.06 resembles the fragment of the N-containing hetero ring structure (C_5_H_8_N^+^, dihydropyridine) after loss of the hydroxyl function and carboxylic acid methyl-ester function. Together, the structure of the infection-dependent NHP-derived compound MeNHP as NHP-methyl-ester was confirmed.

To strengthen the hypothesis that MeNHP is a downstream metabolite of NHP, we investigated the influence of *ald1* loss-of-function mutation on the occurrence of MeNHP *in vivo* (Fig. 2d). We observed that MeNHP accumulated in Col-0 plants 24 hours post UV (hpUV) stress. Col-0 control plants and *ald1* control, as well as, *ald1* UV-treated plants did not show any signal for MeNHP 24 hpUV. Moreover, we tested *FMO1-3D* overexpression lines and *FMO1-3D ald1* double mutant plants on their basal MeNHP amount (Fig. 3D). We observed a constitutive accumulation of MeMHP in the *FMO1-3D* mutant background, whereas no MeNHP was detected in the *FMO1-3D ald1* background.

In conclusion, these data further support that MeNHP is NHP-methyl-ester produced *in planta* downstream of NHP. Mutations in the major biosynthetic genes *ald1* and *fmo1* lead to absence of MeNHP after stress induced biosynthesis. Additionally, we showed that NHPMT1 was able to catalyze the formation of MeNHP from NHP and SAM *in vitro* and therefore, provide additional data that confirm exact mass and retention time information from *in vivo* MeNHP and chemically synthesized authentic standard. Whether MeNHP has an influence on plant-immunity remains to be investigated. It remains to be determined whether additional molecules other than MeNHP, NHP-*O*-Glc and NHP-Glc-ester derive from NHP directly and are present in the Pip/NHP molecular network *in vivo*.

### Identification of NHP-derived metabolites via NHP and D_9_-NHP leaf infiltration, UV-treatment and non-targeted metabolomics

To confirm the occurrence of the observed NHP-derivative after stress application and to screen for additional NHP-derivatives, a non-targeted metabolome experiment was performed as another independent line of evidence. We applied NHP and D_9_-NHP co-infiltration, as well as, mock infiltration with 10 mM MgCl_2_ in leaves of WT, *fmo1-1* and *ugt76b1-1* plants, respectively, and treated the leaves with UV-C afterwards. The aim of this experimental setting was to identify all NHP-derived metabolites, by selecting pairwise features with a mass shift of 9.056 Da (exchange of all 9 hydrogens of the pyridine-moiety by deuterium in NHP) and a small retention time shift of < 0.13 min that are enriched after NHP/D_9_-NHP infiltration, and ideally be synthesized *in vivo*, without external application. In addition, *ugt76b1-1* mutant plants were included as we hypothesized that other NHP-derived metabolites will accumulate as alternative routes for NHP-turnover, when NHP-*O*Glc cannot be synthesized.

The non-targeted metabolite fingerprinting identified 1152 metabolite features with a false discovery rate (FDR) < 10^−5^. The representation of these features by pattern-based clustering via an one-dimensional-self organizing map (1D-SOM) shows two clusters (Fig. 4, cluster number 1 and 2), where metabolites accumulated as consequence of NHP/D_9_-NHP co-infiltration. Cluster 1 shows metabolites accumulating after NHPs infiltration in an *UGT76B1*, as well as, UV-C dependent manner. Here three feature pairs were detected with a mass shift of 9.056 Da. First, the pair NHP-*O*Glc/D_9_-NHP-*O*Glc was detected in cluster 1, as NHP-*O*Glc is known to be exclusively synthesized by UGT76B1 *in vivo*. This chemotype cannot be restored by external application of NHPs to the *ugt76b1-*1 mutant plants. Furthermore, NHP-*O*Glc accumulated also in those Col-0 samples after UV-stress, where NHP/D_9_-NHP infiltration did not take place. As expected, D_9_-NHP-*O*Glc was present in Col-0 and *fmo1-1* plants after NHPs infiltration. The second pair of features with a mass shift of 9.056 Da has exact masses of [M+H]^+^/[D_9_M+H]^+^ 470.185/479.241. The exact mass information and its UGT76B1-dependancy let us tentatively assign the features as NHP-*O*Glc-Hex/D_9_-NHP-*O*Glc-Hex. Subtracting the exact mass of NHP-*O*Glc ([M+H]^+^ 308.134) from the molecule of [M+H]^+^ 470.185 results in a fragment of 162.051 Da, which corresponds to a hexose moiety. Since the feature pair is exclusively detected in the lines with functional UGT76B1, it strongly suggests that UGT76B1 is responsible for the *O*-glycosylation of NHP required for NHP-*O*Glc-Hex synthesis. In-source fragmentation analysis underlines the compound identity by the detection of NHP-*O*Glc as fragment ion of [M+H]^+^ 308.134 (Fig. S3). In addition, NHP-*O*Glc-Hex is present in mock infiltrated UV-stressed Col-0 plants, which confirms NHP-*O*Glc-Hex as a native NHP-derivative. The third pair of features in cluster 1 fits to NHP-*O*Glc/D_9_-NHP-*O*Glc with an additional C_3_H_3_O_3_-moiety, [M+H]^+^/[D_9_M+H]^+^ 394.132/403.188. It is *UGT76B1*-dependent, too, as the molecular features are not present in the *ugt76b1-1* background. A small amount of NHP-*O*Glc-C_3_H_3_O_3_ in mock infiltrated, UV-stressed Col-0 plants confirmed that this metabolite is a native NHP-derivative. MS/MS-fragment analysis of the unknown molecule yielded a fragment of the NHP-backbone of [M+H]^+^ 308.134, and suggests for an additional malonic acid residue of *m/z* 87.007 as the fragment ion (Fig. S4). Together the MS data led us to assign the third feature pair as NHP-*O*Glc-malonic acid.

**Fig. 4.**
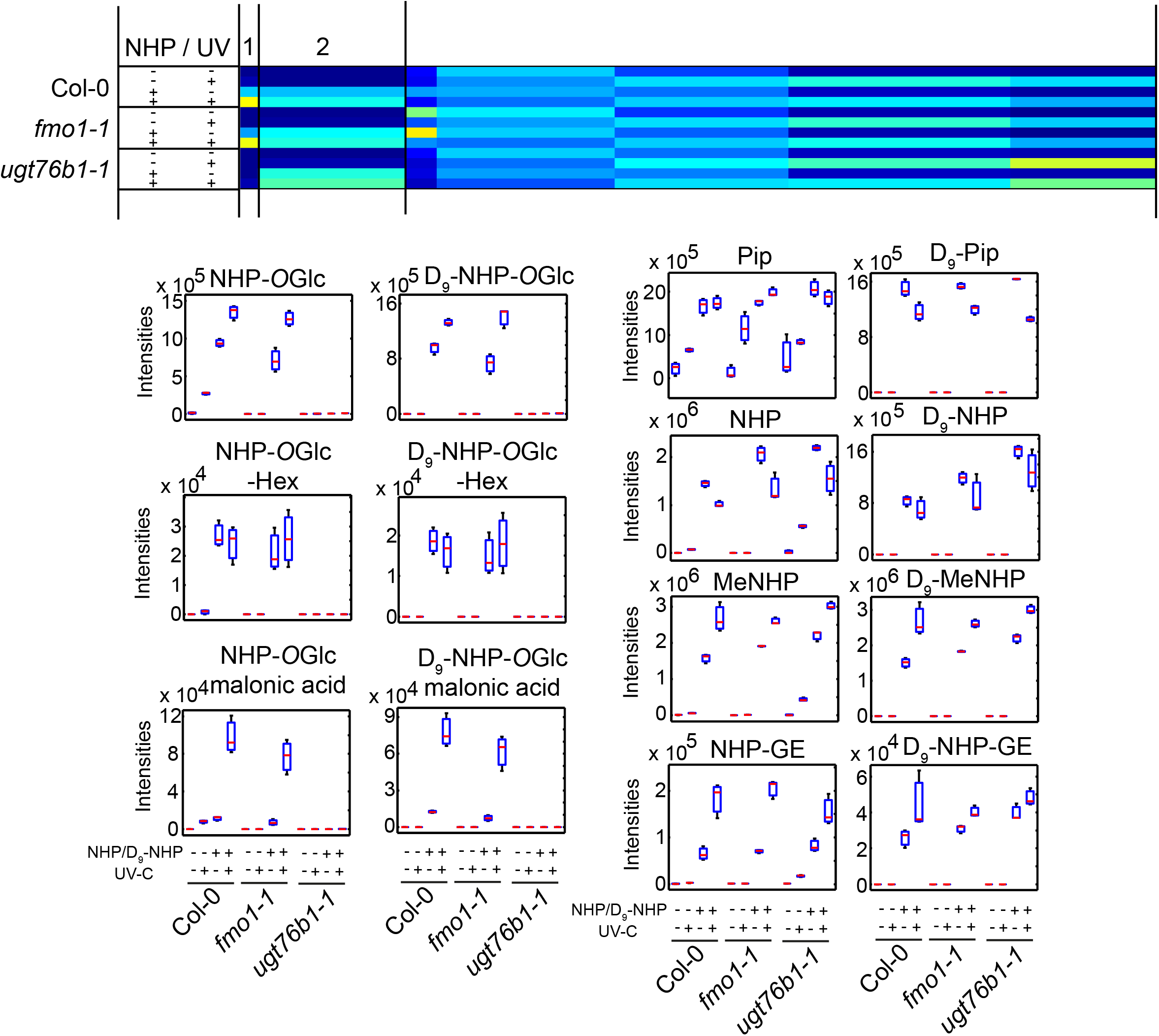
Co-infiltration of NHP/D_9_-NHP into Col-0, *fmo1-1* and *ugt76b1-1* with a subsequent UV-C-trigger underlines additional *in planta* synthesis of NHP-derivates. Into unstressed leaves, either mock solution (10 mM MgCl_2_) or a mixture of 1 mM NHP and 1 mM D_9_-NHP were infiltrated. Plants were kept untreated as control or stressed afterwards for 20 min with UV-C light. The plants were incubated for 24 hours in short day conditions before infiltrated leaves were harvested. The extracted leaf material was analyzed with UHPLC-HRMS. Data was analyzed using Profinder 10.0 (Agilent Technologies, Santa Clara, CA, USA) and MarVis (Kaever et al. 2015. Each sample represent an independent pool of 6 infiltrated leaves of two plants each. Data are shown in Box-Plots representing mean and standard deviation.

Cluster 2 represents metabolites that enrich in all three genotypes after NHP/D_9_-NHP infiltration, with and without UV-C treatment. Via mass shift search, we detected Pip/D_9_-Pip and NHP/D_9_-NHP, MeNHP/D_9_-MeNHP and NHP-GE/D_9_-NHP-GE as NHP-derived metabolites. Pip, NHP and MeNHP accumulated in Col-0 and *ugt76b1-1* after UV-stress. In contrast, NHP-GE was detected only in *ugt76b1-1* after UV-stress in case the NHP/D_9_-NHP mixture was not additional infiltration. Interpretation of the MS/MS-fragment pattern confirmed the identity of NHP-GE (Fig. S5).

Together the experiment expands the number of novel NHP metabolites to MeNHP, NHP-*O*Glc-Hex, NHP-*O*Glc-malonic acid. Furthermore, the conversion of D_9_-NHP into D_9_-Pip led us to propose a so far unknown additional dehydration reaction.

### MeNHP application rescues the susceptibility of fmo1-1 against Hyaloperonospora arabidopsidis Noco 2 infection

To determine the metabolic fate of externally applied MeNHP and to assign its role in plant immunity, 0.1 mM MeNHP were infiltrated into leaves of Col-0, *fmo1-1* and *fmo1-1 ugt76b1* plants. Inspired by that SA-binding protein 2 (SABP2) and some of its Arabidopsis orthologous hydrolyze MeSA to SA, we wondered whether MeNHP is hydrolyzed to NHP in Arabidopsis as well (Forouhar *et al*., 2005; Vlot *et al*., 2008a). In addition, we aimed to figure out, which other NHP-related metabolites accumulate upon infiltration of MeNHP and whether SA biosynthesis is induced by MeNHP or by MeNHP-derived metabolites. Furthermore, the mutant *fmo1 ugt76b1* was included to identify the origin of MeNHP-*O*Glc detected in our initial experiment described above (Fig. S6). After MeNHP-infiltration, MeNHP was detected in all three genotypes (Fig. 5), with a higher signal intensity in *fmo1 ugt76b1*. A comparable intensity pattern was observed for NHP, which was significantly enriched after MeNHP treatment in all three backgrounds and accumulated the most in *fmo1 ugt76b1*, which hints towards hydrolysis of the infiltrated MeNHP. Interestingly, the relative amount of Pip increased significantly in Col-0, *fmo1-1, fmo1 ugt76b1* plants after MeNHP infiltration in comparison to mock treated plants. NHP-*O*Glc was not detected in *fmo1 ugt76b1* plants, but significantly accumulates in Col-0 and *fmo1-1*. NHP-GE accumulates in all three backgrounds. Interestingly we identified a signal of m/z 322.149, which may represent MeNHP-*O*Glc. MeNHP-*O*Glc accumulated significantly after MeNHP infiltration independent of *UGT76B1*. To underline the identification, we conducted an enzymatic reaction using purified SAG forming enzyme UGT74F1 and were able to reproduce the MeNHP-*O*Glc signal *in vitro* (Fig. S6). Furthermore, MeNHP treatment resulted in neither a signal increase of SA nor the accumulation of SAG compared to mock treatment. Similar data were obtained after spraying MeNHP to Col-0 and *fmo1-1* plants (Fig. S7). From these results, we conclude that MeNHP can be metabolized in the plant after external application. We were able to detect accumulation of Pip, NHP, NHP-*O*Glc, NHP-GE and MeNHP-*O*Glc in Col-0 but more important in *fmo1-1* knock-out plants.

**Fig. 5.**
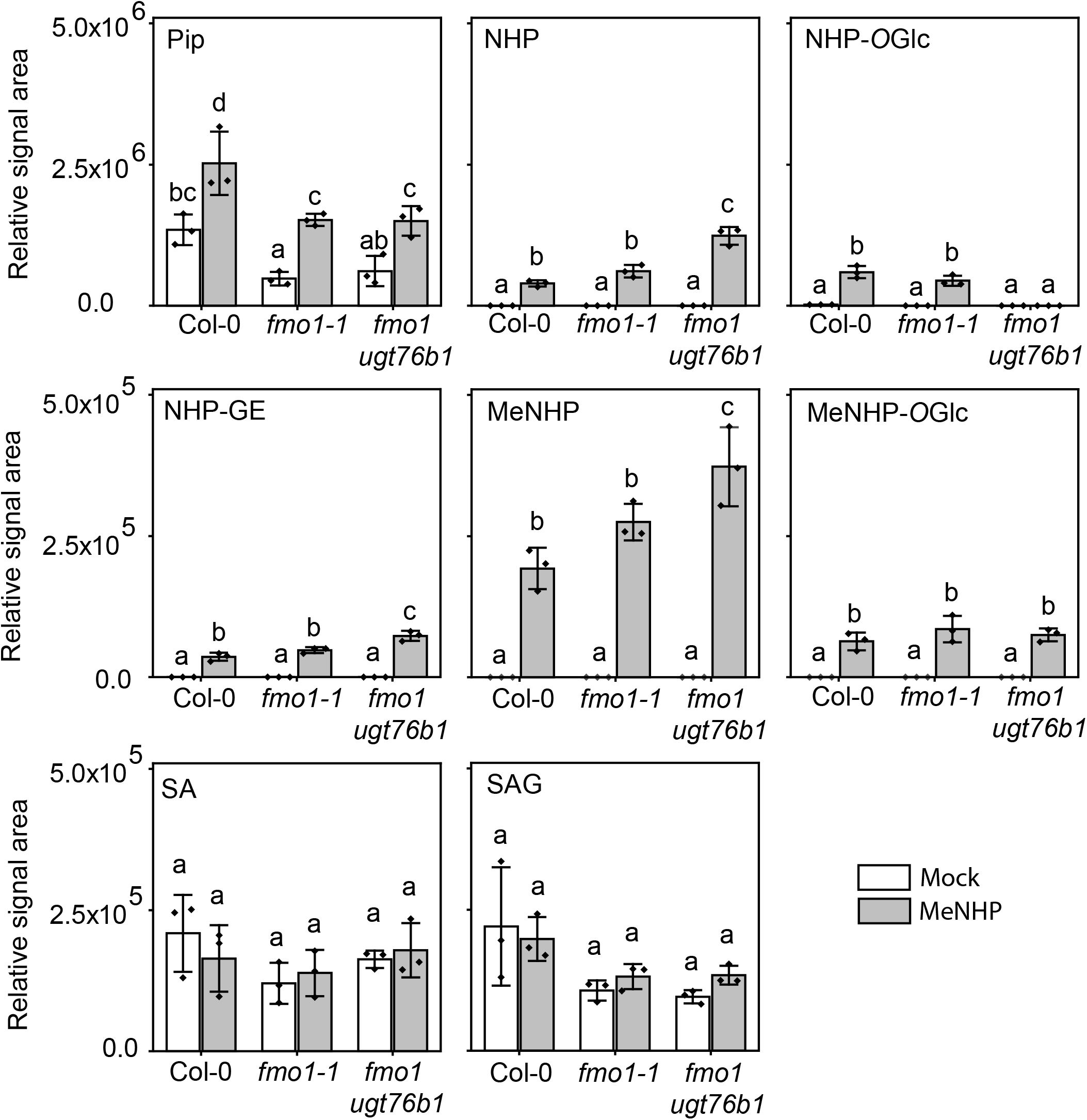
MeNHP-infiltration leads to metabolite remodeling including NHP and Pip accumulation in *fmo1-1* mutant background. Col-0, *fmo1-1* and *fmo1ugt76b1* were grown in short day conditions for 5 weeks. Plants were infiltrated with 10 mM MgCl_2_ (Mock) or 1 mM MeNHP in 10 mM MgCl_2_ (MeNHP). The plants were incubated overnight and harvested 20 hours post infiltration. Samples were extracted with 80 % MeOH and measured with UHPLC-HRMS. Data were analyzed using Qualitative analysis (Agilent Technologies, Santa Clara, CA, USA) for relative signal area of the respective compound shown. Relative signal areas of MeNHP, SA, SAG, Pip, NHP, NHP-*O*Glc, NHP-GE and MeNHP-*O*Glc are shown in mock and MeNHP treated Col-0, *fmo1-1* and *fmo1ugt76b1*. Each replicate represents an independent pool of 6 infiltrated leaves from two plants. Data represent mean with standard deviation. Statistical analysis was done using Origin Pro 2020. Letters indicate statistical differences (p < 0.05, one-way ANOVA post-hoc tukey-test).

To test whether MeNHP is able to prime defense response in Arabidopsis and further to rescue the *fmo1-1* infection phenotype, we challenged MeNHP treated *fmo1-1* and Col-0 plants with a spore solution of *H. arabidopsidis* Noco 2 and analyzed spore growth on mock or MeNHP treated plants (Fig. 6A and B). A concentration gradient of 200 µM, 125 µM, 20 µM and 1 µM was applied to individual groups of *fmo1-1* mutant plants and spore growth was analyzed in comparison to mock treatments (Fig. 6A). We assayed two individual mock treatments against either 200 µM and 125 µM or 20 µM and 1 µM MeNHP. With treatment of 200 µM MeNHP, 69 % of the pathogen growth was assigned to disease category zero (no spore growth), 8 % to category one and 23 % to category two. In the respective mock treatment, growth of the pathogen was grouped into disease category five at 100 %. The comparison shows reduced pathogen sporulation, therefore, lower disease categories with 200 µM MeNHP treatment compared to mock. The 125 µM MeNHP treatment resulted in a similar trend of reduced pathogen growth. Plants pretreated with 125 µM MeNHP exhibited no spore growth at 15 % and disease categories one at 23 %, two at 38 %, three at 8 % and four at 15 %. A comparison between 20 µM MeNHP and mock treatment showed a similar trend as above. Spore growth on mock treated plants was assigned to disease categories four 7 % and five 93 % of plants, whereas spore growth on plants treated with 20 µM MeNHP was grouped into disease categories two 53 %, three 7 %, four 27% and five 13 %. At the lowest concentration of 1 µM MeNHP 10 % of plants group into category three, 20 % in category four and 70 % of the challenged plants group into disease category five. We next applied 200 µM and 125 µM MeNHP to Col-0 and *fmo1-1* plants, respectively, to compare spore growth of *H*.*a*. Noco 2 (Fig. 6B). Mock treated Col-0 plants group into disease categories five and four. Treatment with 200 µM MeNHP resulted in no spore growth on Col-0. Treatment with 125 µM MeNHP resulted in disease categories zero and one. Mock treated *fmo1-1* mutant plants group into disease category five. Treatment with 200 µM MeNHP resulted in categories zero, one and two. Application of 125 µM MeNHP resulted in spore growth that was grouped into disease categories zero, two and three.

**Fig. 6.**
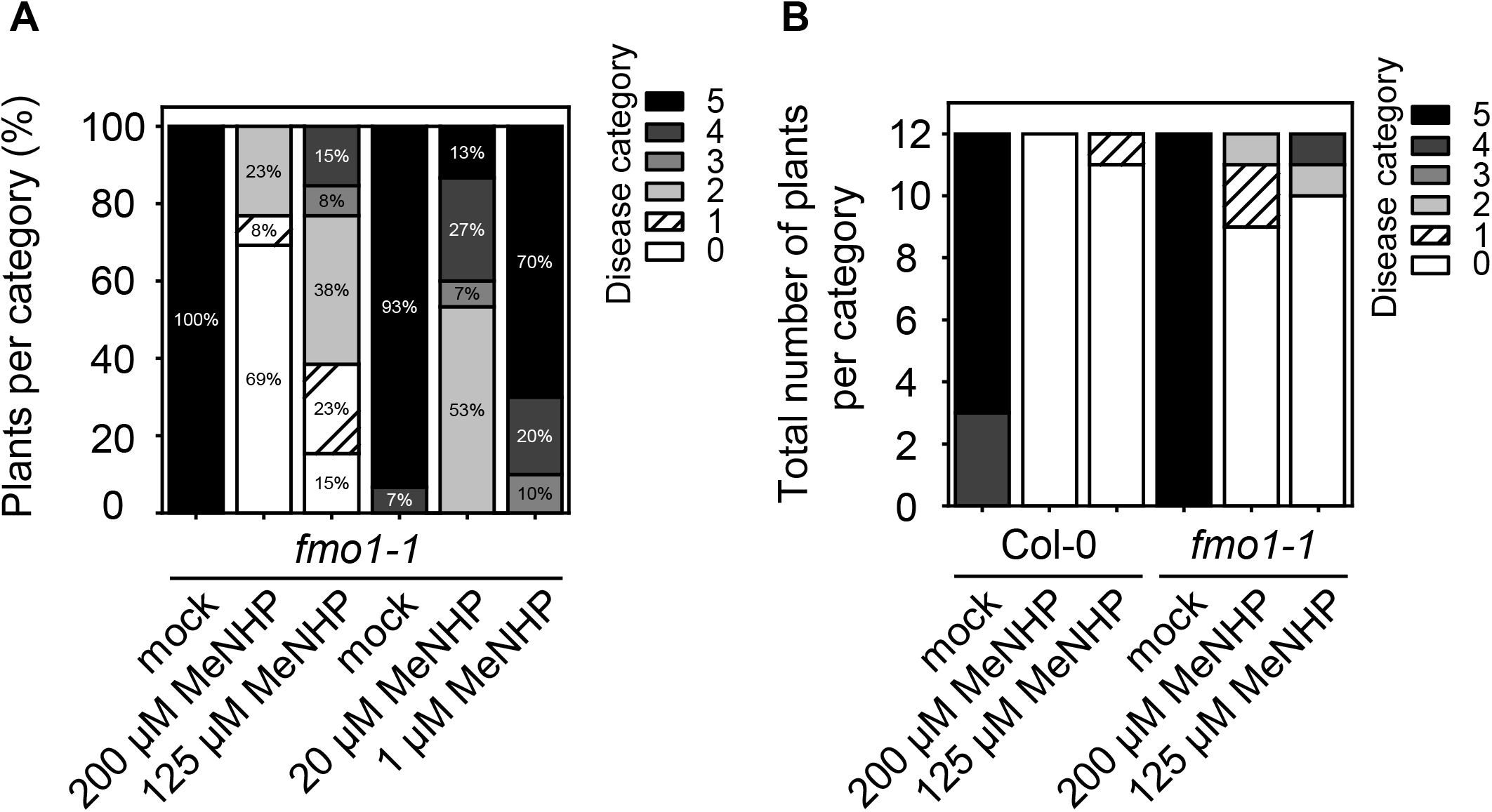
MeNHP infiltration rescues the *fmo1-1* infection phenotype against *Hyaloperonospora arabidopsidis* Noco 2 compared to mock treatment. **A** Four different concentrations of MeNHP (200 µM, 125 µM, 20 µM and 1 µM) were applied to *fmo1-1* mutant plants and *H. arabidopsidis* Noco 2 spore growth was assayed compared to individual mock (water) treated plants. **B** Col-0 and *fmo1-1* mutant plants were treated with MeNHP at a concentration of 200 µM or 125 µM. *H. arabidopsidis* Noco 2 spore growth was assayed compared to individual mock (water) treated plants. Plants were grouped into disease categories from 5 (heavy spore growth) to 0 (no spore growth). Disease categories were assigned as following: Category 5 = more than two leaves harbor > 5 spores, 4 = two leaves harbor > 5 spores, 3 = two leaves < 5 spores, 2 = one leaf > 5 spores, 1 = one leaf < 5 spores, 0 = no spores at all. n= 10-15 plants per treatment.

The experiments confirm that the applied MeNHP is metabolized to NHP in Arabidopsis and that MeNHP treatment is able to rescue the susceptible phenotype of NHP biosynthesis mutant *fmo1-1* and to induce resistance in Col-0. Furthermore, MeNHP treatment alone does not lead to an increase in signal intensity of SA and SAG, however, the amounts of Pip, NHP, NHP-*O*Glc, NHP-GE and MeNHP-*O*Glc increase significantly.

## Discussion

Intact metabolite networks are key to hormonal balance in plants. In this work, we lay out the NHP-metabolome by non-targeted UHPLC-HRMS-based metabolomics. For the initial approach, a non-targeted dataset of Arabidopsis infection with *Pseudomonas* was recorded. The strategy was to identify NHP-metabolite features based on exact mass information of the sum formula. *In silico* modifications were performed, based on well-known metabolizing reactions, such as hydroxylation, methylation and amino acid conjugation. From the sum formula of the designed compound, its exact mass was identified and molecular identification was targeted. Via both, analysis of *P*.*s*.*m*. infiltrated Col-0 and *fmo1-1* mutant plants and dual infiltration of NHP/D_9_-NHP, three molecules of NHP-turnover were identified, namely, MeNHP, NHP-*O*Glc-Hex and NHP-*O*Glc-malonic acid (Fig. 2, 3 and 4). We detected all three metabolites in Col-0, however, not in the *fmo1-1* mutant background after *P*.*s*.*m*. infiltration or UV-C treatment. Moreover, we were able to show that MeNHP is metabolized to NHP and that MeNHP treatment is able to rescue the susceptible phenotype of *fmo1-1* mutant plants against *H. arabidopsidis* Noco 2.

### Dual-infiltration as unbiased method to detect undescribed metabolites of NHP

The dual-infiltration method was developed to overcome detection limitation of minor metabolites from native plant extracts and to enable unbiased molecular feature identification, independent of a targeted screen. Due to adding both authentic standard and D_9_-labeled authentic standard, the sensitivity to detect NHP metabolites was increased. Especially, the specificity to pin down a molecule to be of NHP origin was enhanced. By the distinctive mass shift fingerprint and retention time difference, we are able to assign metabolites to NHP-origin. Together with the possibility to identify the metabolites in the UV-stressed Col-0 plants, the analysis gives a broad picture of the NHP-metabolites. Most importantly, we are able to present molecules absent in the *fmo1-1* background underlining functional *FMO1*- and NHP-dependency. The data acquisition and analysis enclose both ionization modes positive ESI and negative ESI. To ensure high quality features, molecules with FDR < 10^−5^ are included into the dataset underlying non-targeted analysis of D_9_-labled and unlabeled molecular pairs. Yu and colleagues published a similar labeling approach to describe the ability of different species to metabolize nematode signaling molecules (Yu *et al*., 2021). The researchers applied ascarosides and C^13^-labeled ascarosides to several organisms to identify their ability to metabolize the Nematode derived compounds by product and C^13^-labeledeld product analysis. Similarly, the group chose exact mass and retention time shift as quality measure for unbiased non-targeted analysis (Yu *et al*., 2021). The successful application including metabolic turnover inspired us to investigate metabolite mass shifts with our NHP and D_9_-labeled NHP standard. NHP metabolites that have already been described are two glycoside forms NHP-*O*Glc and NHP-GE (Bauer *et al*., 2021; Chen *et al*., 2018; Hartmann *et al*., 2018). Whereas the biosynthesis and infection dependency of NHP-*O*Glc have been characterized independently, the unambiguous identification of NHP-GE needs to be underlined and its route of biosynthesis remains unknown (Bauer *et al*., 2021; Cai *et al*., 2021; Holmes *et al*., 2021; Mohnike *et al*., 2021). We tested activity of heterologous expressed and purified UGT73D1 against NHP *in vitro*, however, were no NHP-GE synthesizing activity was found (Fig S8.). In our analysis, NHP-GE is favorably detected in the *ugt76b1* background but of very low to no abundance in Col-0 plants after *P*.*s*.*m*. or UV treatment. As proof of the dual-infiltration concept, we showed the expected molecular feature pairs of NHP/D_9_-NHP and NHP-*O*Glc/D_9_-NHP-*O*Glc (Fig. 5). NHP and NHP-*O*Glc are missing in the *fmo1-1* mutant background without external application of NHP/D_9_NHP, but are present when treated with the mixture. Similarly, the MeNHP signal was missing in the *fmo1-1* background and dual-infiltration restored the MeNHP/D_9_-MeNHP signal, respectively. Additionally, we were able to detect *UGT76B1*-dependent NHP-metabolites, namely, NHP-*O*Glc-Hex and NHP-*O*Glc-Mal. To collect evidence for their molecular structure, we analyzed mass spectra of the two compounds. Infiltrated plant metabolite extract was subject to spectra analysis. In source fragment ions, underline the identification of NHP-*O*Glc-Hex as the insource fragment *m/z* 308.134 represent NHP-*O*Glc (Fig. S3). Fragment spectrum analysis suggests malonic acid addition to NHP-*O*Glc represented by a fragment ion at *m/z* 87.008 (Fig. S4). Malonic acid moieties at glucose residues are for example present at anthocyanin’s (Bloor and Abrahams, 2002). Interestingly both molecules are synthesized *in vivo* after UV-stress without the need of additional infiltration.

### Structure elucidation and NHP-dependency of MeNHP synthesis

The discovery of the molecular feature *m/z* 160.097, which was underlined by the pairwise feature identification in dual-infiltration suggested NHP methylation. Nevertheless, we could only rely on the predicted exact mass information of the expected molecular formula that resembles MeNHP. NHP-dependency and the site of methylation remained unclear. To underline MeNHP detection and to identify its site of methylation, we chemically synthesized NHP-methyl-ester from pipecolic acid methyl ester. Due to the specificity of methylation at the carboxylic acid function within the MeNHP-standard, we were able to exclude hydroxyl methylation. We underline MeNHP as NHP-methyl-ester via retention time and MSMS-fragment comparison between authentic standard and the *in vivo* signal (Fig. 3c). Both the MeNHP-standard and the *in vivo* signal exhibit a retention time of 2.57 min, as well as similar fragmentation behavior. Their fragment ions *m/z* 142.08, *m/z* 127.063, *m/z* 110.060, *m/z* 100.076 and *m/z* 82.065 are identical and derive from the mother mass *m/*z 160.097. Especially the fragment ions *m/*z 127.063, *m/*z 100.076 and *m/*z 82.065 are identical with NHP-fragments, underlining structural similarities and hint towards a NHP-derived molecule (Chen *et al*., 2018; Hartmann *et al*., 2018). To strengthen the hypothesis that MeNHP is a NHP-derived metabolite, we analyzed functional dependency on NHP-biosynthesis. Via analysis of UV-stressed Col-0 against *ald1* or the basal accumulation of MeNHP in *FMO1-3D* against *FMO1-3D ald1* mutant plants, the dependency of MeNHP on functional NHP biosynthesis was stressed further (Fig. 3d). To support exact mass accuracy and retention time data of MeNHP, we present an *in vitro* reaction of NHPMT1, which produced MeNHP by using NHP as substrate and SAM as co-substrate. Both *in vivo* and *in vitro* MeNHP compounds behave as authentic standard in respect to RT and fragmentation pattern. Despite the *in vitro* activity of NHPMT1 with NHP, SALK_053006 (*nhpmt1-1*), SALKseq_135601 (*nhpmt1-2*) and *nhpmt1-1 ugt76b1-1* mutant plants did not show absence of MeNHP signal after UV-treatment. Surprisingly, the signal intensity of NHP and MeNHP was increased in the analyzed mutants (Fig. S9, Fig. S10). The analysis of *nhpmt1* mutant plants raises the question, if redundant MTases exist for MeNHP synthesis, or if NHPMT1 shows promiscuous MTase activity with NHP but has no influence on *in vivo* synthesis.

### Physiological implications of NHP-metabolites

By targeted and non-targeted metabolomics approaches, NHP metabolites were investigated and novel candidate molecules are presented. Additionally, we underline the discovery of NHP-GE by Bauer et al. and present three novel metabolites which are most likely NHP derived and unambiguously *FMO1*-dependent after *P*.*s*.*m*. infiltration. Independently we present the NHP-metabolites in a dual infiltration study, tracking the metabolic fate via non-targeted UHPLC-HRMS metabolomics. However, their physiological implications remain elusive. In contrast to the data present by Bauer and colleagues, we detected accumulation of NHP-GE in *ugt76b1* background (Bauer *et al*., 2021). Furthermore, we describe increased levels of MeNHP in *ugt76b1-1* mutant plants. Taken together we suggest the carboxy methylation and carboxy glycosylation of NHP as alternative route of NHP-turnover, when *O*-glycosylation is not available.

Nevertheless, the volatile nature of the methylated phytohormones MeJA and MeSA draw our attention on MeNHP’s potential to enhance resistance. In analogy to MeSA’s ability to induce systemic resistance in tobacco, we investigated the ability of MeNHP to rescue the *fmo1-1* susceptible phenotype towards oomycete pathogen (Hartmann *et al*., 2018; Park *et al*., 2007). MeSA is proposed to be cleaved by tobacco SABP2 resulting in SA and induced acquired resistance (Forouhar *et al*., 2005; Park *et al*., 2007). Several methylesterases (MES) are present in Arabidopsis that show sequence similarity to the tobacco SABP2, of which MES-1, -2, -4, -7 and -9 exhibit *in vitro* activity with MeSA in competition with SA (Vlot *et al*., 2008b). The molecular structure of SA and NHP opens the question, if there is a shared MES capable to hydrolyze MeSA and MeNHP in Arabidopsis, similar to their shared mechanism for glucosylation by UGT76B1 (Mohnike *et al*., 2021; Zeier, 2021). In Figure 5, we lay out NHP-related metabolites accumulated upon infiltration of MeNHP. The data suggest a hydrolysis of the externally applied MeNHP to NHP. The external application of MeNHP did not result in significant changes to the SA levels, neither in infiltration studies, nor after spray application (Fig. 5, Fig. S7). Afterwards, we infiltrated MeNHP into *fmo1-1* mutant plants to investigate the potential to enhance disease resistance, especially aiming to rescue the susceptible phenotype of the *fmo1-1* mutants. We analyzed the spore count of *H*.*a*. Noco 2 on Arabidopsis leaves pretreated with mock or various concentrations of MeNHP (Fig. 6A, B). The data suggests that MeNHP treatment is able to rescues the susceptible phenotype of *fmo1-1* mutant plants, resulting in reduced spore growth. The enhanced resistance after MeNHP treatment at different concentration might be due to the successful conversion of MeNHP to NHP in the *fmo1-1* background. Furthermore, restoring the NHP pool could be a crucial step to enhance disease resistance in the susceptible *fmo1-1* mutant background. In addition, MeNHP treatment increased Col-0 resistance against *H*.*a* Noco 2, too. The applied concentrations ranging from 200 µM to 1 µM used for infiltration are within the range of similar studies that infiltrated NHP to induce defense from 1 mM to 1 µM (Chen *et al*., 2018; Hartmann *et al*., 2018). Additionally, NHP induces SAR in Arabidopsis in low doses independent from the mode of application (Schnake *et al*., 2020). Two independent studies underlined the potential of NHP to induce resistance beyond the scope of Arabidopsis (Holmes *et al*., 2019; Schnake *et al*., 2020). The successful induction of resistance after application of methylated compounds like MeSA and MeJA puts MeNHP in scope for future research in plant-to-plant communication. We suggest a sender receiver experiment applying stress to WT and *fmo1* mutant plants as sender and analyze the NHP chemotype of unstressed *fmo1* receiver plants. In the ideal case, the experiment would be conducted with a MeNHP synthesis mutant.

Interestingly we were able to identify another metabolite of MeNHP, when tracking its metabolic fate, namely, MeNHP-*O*Hex. This compound is *UGT76B1* independent, suggesting an UGT able to use MeNHP as substrate, conjugating the glucosylation at the *N-*hydroxyl function. *In vitro* reaction using UGT74F1 resulted in reproduction of the MeNHP-*O*Glc signal (Fig. 7, Fig. S3). UGT74F1 might be another candidate protein for *in vivo* biosynthesis of the NHP-metabolite. Nevertheless, MeNHP-*O*Glc was not shown to be a native product in plant stress response without external application of MeNHP. Hypothetically, UGT71C3 capable of synthesizing MeSA-*O*Glc may be another interesting candidate protein, due to the similarity in structure between MeNHP and MeSA (Chen *et al*., 2019). We describe that Pip might also be a product of NHP-turnover, as not only Pip was accumulating after the dual infiltration of NHP and D_9_-NHP but also D_9_-Pip. It raises the question for a reaction to remove the *N*-hydroxylation from NHP via, for example, hydrolases or as FMO1-reverse reaction. The observation of NHP cleavage may also explain why Pip amounts are still increasing in time-course experiments, when the NHP, NHP-*O*Glc are already decreasing in signal intensity (Hartmann and Zeier, 2019).

**Fig. 7.**
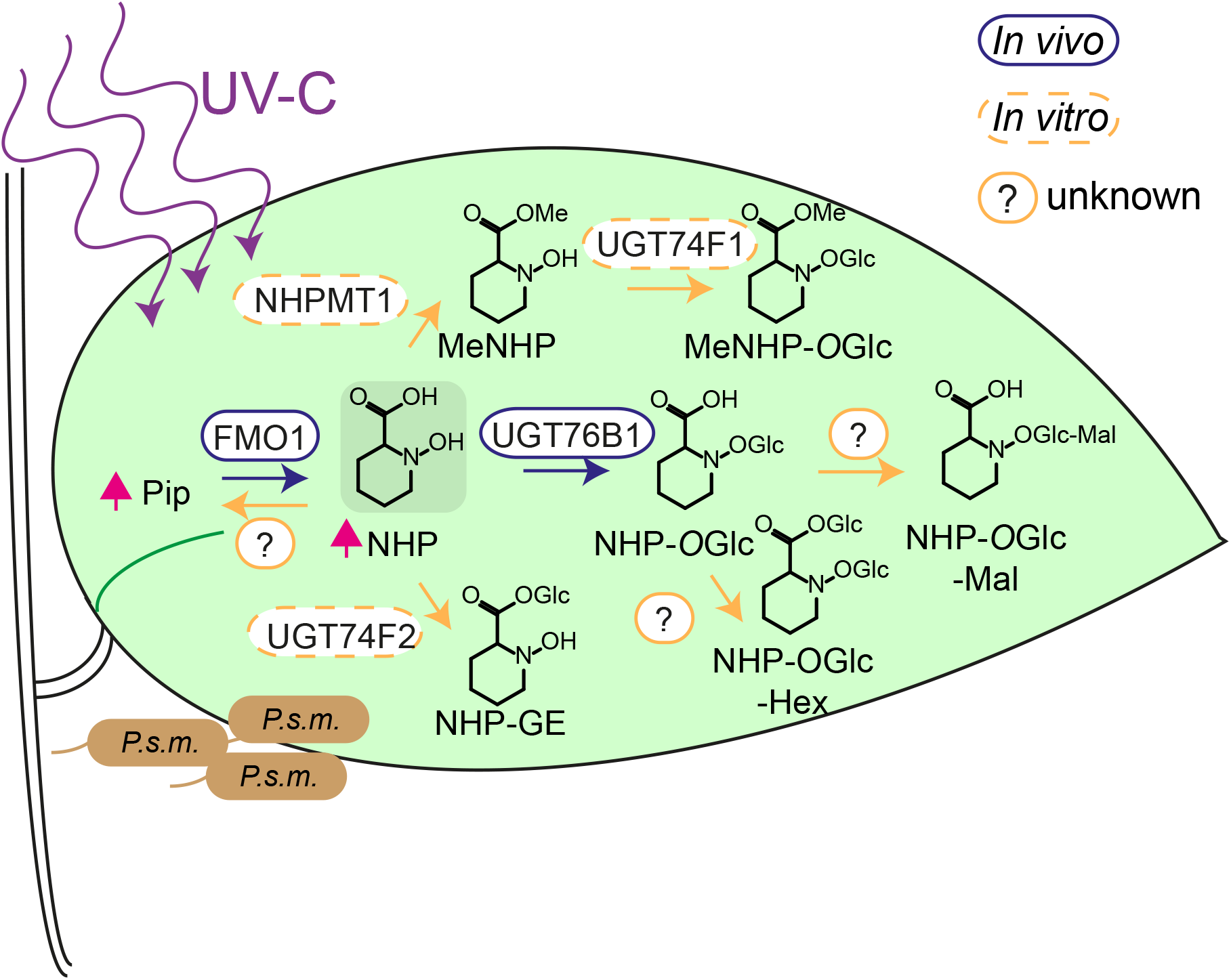
NHP turnover in Arabidopsis induced by UV-C and *P*.*s*.*m*. infiltration. The scheme concludes the detected metabolic routes of NHP-turnover. In response to UV-C stress and *P*.*s*.*m*. infection, Arabidopsis synthesizes Pip and NHP. To reduce cellular NHP levels, it is hydrolyzed, glycosylated and methylated. Respective products were shown to be further metabolized to complex conjugates, NHP-*O*Glc-Hex and putatively to NHP-*O*Glc-malonic acid. Identified enzymes from *in vivo* are circled in purple. Enzymes with identified *in vitro* activity are circled in yellow. Reactions with neither enzyme described *in vivo* nor *in vitro* are marked with a question mark.

An additional mechanism to control hormonal activity is amino acid conjugation. On the one hand, conjugation can lead to activation as it is known in the case of JA and isoleucine by the enzyme JASMONATE RESPONSE LOCUS 1 (JAR1). The JA-modification by JAR1 results in the biological active form JA-Ile (Staswick and Tiryaki, 2004; Suza and Staswick, 2008). On the other hand, inactivation can be achieved. One example is the conjugation of aspartic acid (Asp) to SA in *A. thaliana* by GH3.5. The product SA-Asp is supposed to be biological inactive and a storage metabolite of SA (Chen 2013). Following the pairwise analysis of labeled and unlabeled metabolites, we can exclude the occurrence of NHP-amino acid conjugates in our experimental setting.

## Conclusion

Four novel metabolites were identified via UHPLC-HRMS: MeNHP, MeNHP-*O*Glc, NHP-*O*Glc-Hex and NHP-*O*Glc-Mal (Fig. 7). The potential of MeNHP to induce defense priming was investigated. Further research, however, is required to clarify the role of MeNHP in defense response, for example, in plant-to-plant communication. What is more is that metabolites of NHP accumulate in *ugt76b1* mutants, where an important mode of NHP turnover into NHP-*O*Glc is unavailable, suggesting the ability to shuttle NHP into other metabolic pathways to certain extent.

## Supplementary data

Fig. S1. SDS-PAGE from ion metal affinity chromatography purification of heterologously expressed AT4G22530 (NHPMT1).

Fig. S2. Collision induced dissociation fragments of MeNHP and NHP

Fig. S3. Insource fragments of NHP-*O*Glc-Hex and D_9_-NHP-*O*Glc-Hex feature pair.

Fig. S4. Collision induced dissociation fragments of NHP-*O*Glc-malonic acid and D_9_-NHP-*O*Glc malonic acid feature pair.

Fig. S5. Collision induced dissociation fragments of NHP-GE.

Fig. S6. Infiltration of MeNHP leads to MeNHP-*O*Glc formation, which is underlined *in vitro*.

Fig. S7. Metabolite analysis after spray application of MeNHP.

Fig. S8. UGT73D1 is not active with NHP *in vitro*.

Fig. S9. Metabolite analysis of UV-treated Col-0 and *nhpmt1* mutant plants.

Fig. S10. MeNHP analysis of UV-stressed Col-0 vs. *ugt76b1-1 nhpmt1-1*.

Supplemental Protocol S11. Chemical synthesis of MeNHP.

Supplemental Dataset S12. UHPLC-HRMS-data of NHP/D9NHP dual-infiltration.

## Acknowledgements

We like to thank Prof. Dr. Christiane Gatz for fruitful discussion and Prof. Dr. Armin Djamei for proving the 80 % MeOH extraction protocol. We thank Dr. Ellen Hornung for cloning of NHPMT1 and crossing of *ugt76b1-1* with *nhpmt1-1* mutant plants to generate the double mutant. We acknowledge Moritz Klein and Sönke Beewen for laboratory support.

## Author contributions

L.M., W.H., B.W., K.F., Y.Z. and I.F. conceived and designed the experiments. L.M., W.H., B.W. and K.F. performed the experiments. L.M., W.H., B.W., K.F., Y.Z. and I.F. analyzed and discussed the data, L.M., W.H., K.F., Y.Z. and I.F. wrote the article.

## Data availability

All data generated or analyzed during this study are included in this published article and its supplementary information files.

## Abbreviations

1D-SOM: one-dimensional-self organizing map
ALD1: AGD2-LIKE DEFENSE RESPONSE PROTEIN 1
BSMT1: BENZOIC ACID/SA METHYL-TRANSFERASE 1
CRISPR: Clustered Regularly Interspaced Short Palindromic Repeats
DHBA: di-hydroxy benzoic acid
EPS1: ENHANCED PSEUDOMONAS SUSCEPTIBILITY 1
ESI: electrospray ionization
hpUV: hours post ultra-violet light treatment
ICS1: ISOCHORISMIC ACID SYNTHASE 1
FMO1: FLAVINE-DEPENDENT MONOOXYGENASE 1
MeNHP: *N*-hydroxy pipecolic acid methyl ester
MeNHP-*O*Glc: MeNHP glycoside
MES: Methyl esterase
MeSA: Salicylic acid methyl ester
MeSAGlc: Salicylic acid methyl ester glycoside
MS: mass spectrometry
MTBE: Methyl-*tert*-butyl ether
NHP: *N*-hydroxy pipecolic acid
NHP-*O*Glc: NHP glucoside
NHP-*O*Glc-Hex: NHP glucoside hexose
NHP-GE: NHP glycosyl ester
NHPMT: NHP methyl transferase
PBS3: AvrPphB SUSCEPTIBLE 3
Pip: Pipecolic acid
*P*.*s*.*m*.: *Pseudomonas syringae* ES4326
RT: retention time
SA: Salicylic acid
S3H: SA-3-hydroxylase
S5H: SA-5-hydroxylase
SABP2: SA-binding protein 2
SAG: Salicylic acid glucoside
SARD4: SYSTEMIC ACQUIRED RESISTANCE DEFICIENT 4
SGE: Salicylic acid glucoside ester
UGT: UDP-dependent glycosyl transferase
UNK: unknown metabolite
UHPLC-HRMS: ultra-high-performance-liquid-chromatography high-resolution mass spectrometry

